# Methods for discovering genomic loci exhibiting complex patterns of differential methylation

**DOI:** 10.1101/021436

**Authors:** Thomas J Hardcastle

**Affiliations:** Department of Plant Sciences, University of Cambridge, Downing Street, CB2 3EA Cambridge, UK

**Keywords:** segmentSeq, baySeq, Methylation, Differential methylation, DCL-dependent methylation, DMR

## Abstract

Cytosine methylation is widespread in most eukaryotic genomes and is known to play a substantial role in various regulatory pathways. Unmethylated cytosines may be converted to uracil through the addition of sodium bisulphite, allowing genome-wide quantification of cytosine methylation via high-throughput sequencing. The data thus acquired allows the discovery of methylation ‘loci’; contiguous regions of methylation consistently methylated across biological replicates. The mapping of these loci allows for associations with other genomic factors to be identified, and for analyses of differential methylation to take place.

The segmentSeq **R** package is extended to identify methylation loci from high-throughput sequencing data from multiple experimental conditions. A statistical model is then developed that accounts for biological replication and variable rates of non-conversion of cytosines in each sample to compute posterior likelihoods of methylation at each locus within an empirical Bayesian framework. The same model is used as a basis for analysis of differential methylation between multiple experimental conditions with the baySeq **R** package. We demonstrate this method through an analysis of data derived from Dicer-like mutants in *Arabidopsis* that reveals complex interactions between the different Dicer-like mutants and their methylation pathways. We also show in simulation studies that this approach can be significantly more powerful in the detection of differential methylation than existing methods.

## Background

Cytosine methylation, foundin most eukaryotes and playing a key role in gene regulation and epigenetic effects[1–3], can be investigated at a genome wide level through high-throughput sequencing of bisulphite treated DNA [4]. Treatment of denatured DNA with sodium bisulphite deaminates unmethylated cytosines into uracil; sequencing this treated DNA thus allows, in principle, not only the identification of every methylated cytosine but an assessment of the proportion of cells in which the cytosine is methylated. Moreover, by comparing these quantitative methylation data across experimental conditions, genomic regions displaying differential methylation can be detected.

The data available for methylation locus finding from bisulphite treated DNA are generated from a set of sequencing libraries. Each library consists of a set of sequenced reads which can be aligned and summarised [5] to report at each cytosine the number of sequenced reads in which the cytosine is methylated, and the number in which the cytosine is unmethylated [6]. Several methods have been proposed to detect differential methylation at the cytosine level, and to identify contiguous differentially methylated cytosines defined as differentially methylated regions (DMRS) [7]. However, these approaches, in not identifying non-differentially methylated regions and unmethylated regions, preclude many strategies for downstream identification of biological significance of differential methylation.

We propose here a new method for methylation analysis based on the notion of methylation ‘loci’; genomic regions defined by the presence of contiguous cytosines whose methylation is correlated across experimental conditions. These loci may thus be assumed to share biogenesis and functional properties [8]. Furthermore, the identification of methylation ‘loci’ and the quantification of methylation within a locus increases statistical power to detect differential methylation. We show that by taking a novel approach in which methylation loci are identified and subsequently analysed for patterns of differential behaviour, we are able to achieve high levels of accuracy in identifying differential methylation. The empirical Bayesian methods employed also allow analysis of multiple patterns of differential methylation to be identified within complex data sets, allowing for detailed downstream analysis of biological mechanisms. We achieve this by adapting our previously described methods for defining siRNA loci from high-throughput sequencing of sRNAs [9], and for a generalised analysis of high-throughput sequencing data [10]. These methods allow for an analysis of differential behaviour in the methylome that accounts both for biological variation between replicates and systemic differences between samples caused by variations in the conversion rates of bisulphite treatment.

We demonstrate these methods in an analysis of methylation in all contexts in mutants of the Dicer-like (DCL) proteins in *Arabidopsis* [11]. In higher plants, Dicer or Dicer-like proteins form a small gene family of sometimes overlapping function in the biogenesis of small RNAs [12]. In *Arabidopsis*, four different DCL proteins exist, acting in a partially redundant manner [13]. Predominantly, DCL1 is involved in the production of 21-nt miRNAs. DCL2 and DCL4 act redundantly and perhaps hierarchically to produce 22 and 21-nt siRNAs. DCL3 produces 24-nt siRNAs, previously identified as the key component of RNA-directed DNA methylation (RdDM). Recent work [14–16] has however emphasised the importance of 21 and 22-nt siRNAs in regulating methylation at at least some loci. By applying the methods developed here we are able to identify multiple patterns of differential behaviour between the Dicer-like mutants and the loci which correspond to these.

We further demonstrate that this approach, in addition to allowing greater flexibility in analysis, offers better performance than a number of existing methods for the analysis of cytosine methylation data by considering a set of comparisons on simulation studies based on WGBSSuite [17], a recently developed stochastic method for generating simulated single base resolution DNA methylation data. We demonstrate high performance under a variety of simulation conditions, with substantial improvements over existing methods in the analysis of small changes in methylation between experimental conditions and for low numbers of biological replicates.

## Results

We develop here a novel approach to analysis of cytosine methylation based on the well-established and widely used open-source R [18] packages segmentSeq [9] and baySeq [19], available on Bioconductor (http://www.bioconductor.org) [20]. These methods allow for the detection of methylation loci from replicated samples in multiple experimental conditions. Where non-conversion rates are estimable these can be incorporated into the distributional assumptions. Following the detection of methylation loci, we are able to use the same distributional assumptions to identify differential methylation both in pairwise comparisons and more complex experimental designs.

### Analysis of the methylome in *dcl* mutants of *Arabidopsis*

We demonstrate the value of this approach in a reanalysis of the methylome in the Dicer-like mutants from the Stroud et al [11] dataset. We identify in a single analysis methylation loci in the *dcl2*, *dcl3*, *dcl4*, *dcl2/4* and *dcl2/3/4* mutants, together with wild-type samples and discover complex patterns of differential methylation that exist between these mutants and the wild-type samples.

We begin with a standard pipeline for read alignment and summarisation [5]. Reads were aligned and summarised for each methylation context using the Bismark caller [21] with default settings. Since cytosine methylation should be absent in the chloroplast and mitochondrial genomes, we can estimate non-conversion rates as the ratio of sequenced cytosines to thymines in the reads aligning to these genomes. This was done for each sample and incorporated into the analysis at the distributional level (see Methods).

We separate the data into the three major contexts of methylation; CpG, CHG, and CHH. For each context of methylation, we identify a set of loci and estimate posterior likelihoods that any given locus is truly methylated in each of the experimental conditions (see Methods). Figure 1 summarises the input data and expected numbers of loci in each mutant, based on the posterior likelihoods. The genome-wide trends in methylation remain relatively constant in all of the dcl mutants relative to the wild-type samples, with some minor loss of methylation (relative to wild-type) at this scale in the CHH context in the dc*l2/3/4* and *dcl3* mutants, and some gain of CHH methylation in *dcl4* mutant at the centromeric regions. The total number of methylation loci in each condition may be estimated by summing the posterior liklelihoods of loci (Fig. 1d). Relative to wild-type, expected numbers of loci do not alter substantially for *dcl2/4* loci in any condition, or for CpG methylation in *dcl2/3/4*, while all the single mutants show lower numbers of methylation in all contexts. The numbers of methylation loci discovered in the CHG context are substantially lower than for other contexts; however, the loci discovered are generally longer, as shown by the estimated portion of the genome covered by loci in each context (Fig. 1e), which shows roughly equivalent coverage for CpG and CHG with a minor reduction in CHH context.

**Figure 1.**
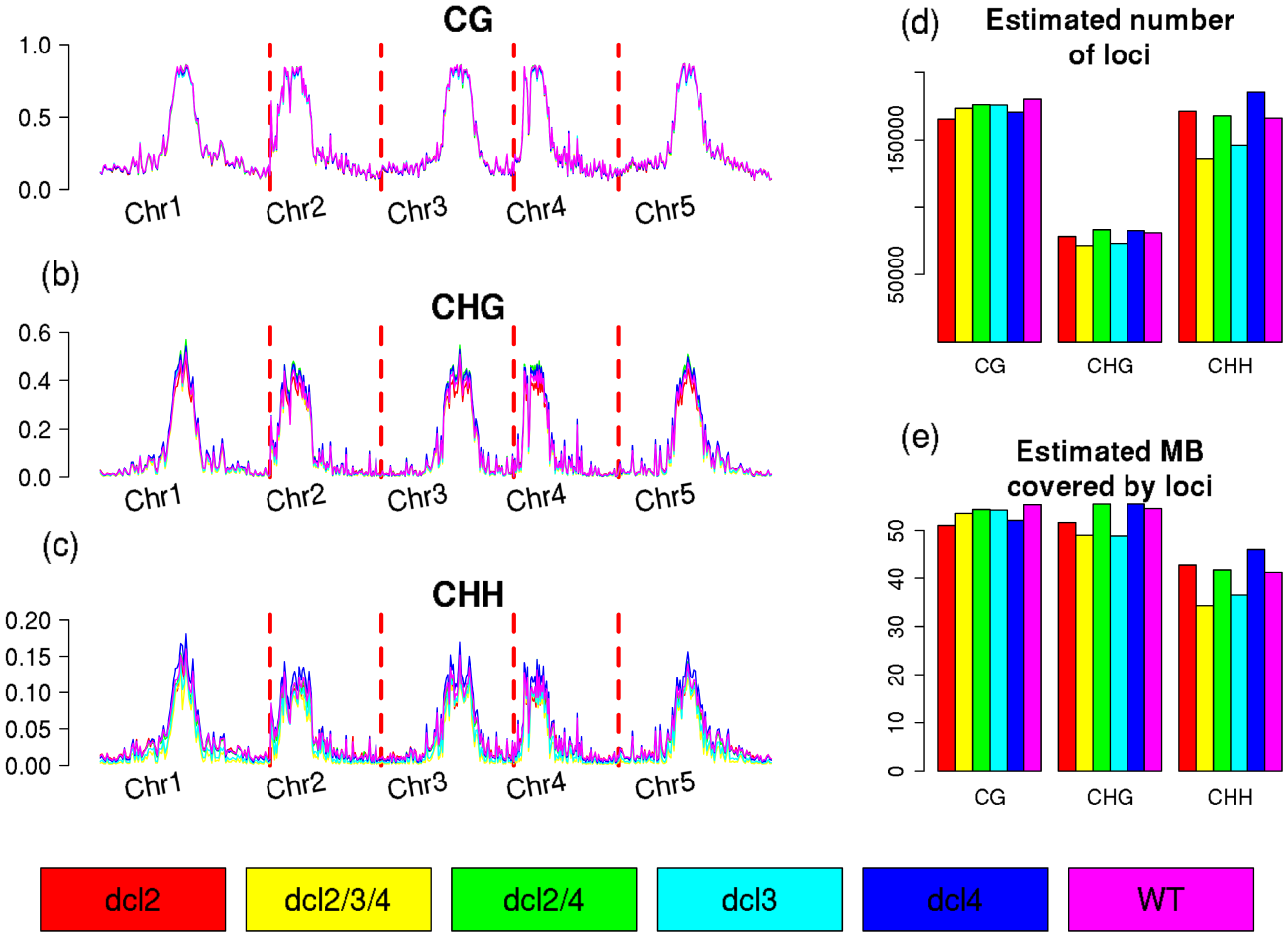
Genome-wide profiles of methylation and expected numbers of loci in dcl mutants. Genome wide profiles of methylation for the various Dicer-like mutants, and wild-type, in CpG (a) CHG (b) and CHH (c) contexts, adjusted for non-conversion rates. The estimated number of loci identified in each condition are shown in (d), while the estimated length of the genome covered by loci are shown in (e).

We next consider patterns of differential methylation at the level of the identified loci. For each region of the genome, posterior likelihoods of difference are identified, and adjusted by the likelihood that the region is a methylation locus in at least one condition. From these posterior likelihoods, we can estimate the expected number of loci belonging to each model of equivalence and difference between the conditions as the sum of the posterior likelihoods for this model over all loci. We can also select specific loci by controlling the false discovery rate (FDR) estimated from the posterior likelihoods. Ten patterns of differential methylation (Figure 2) are identified with an estimated number of loci greater than one thousand and a number of loci with an FDR < 0.05 greater than two hundred in at least one methylation context. Numbers for these models are described in Table 1 while a fuller list of potential models and the numbers of loci corresponding to these is available in Supplementary Materials.

**Figure 2.**
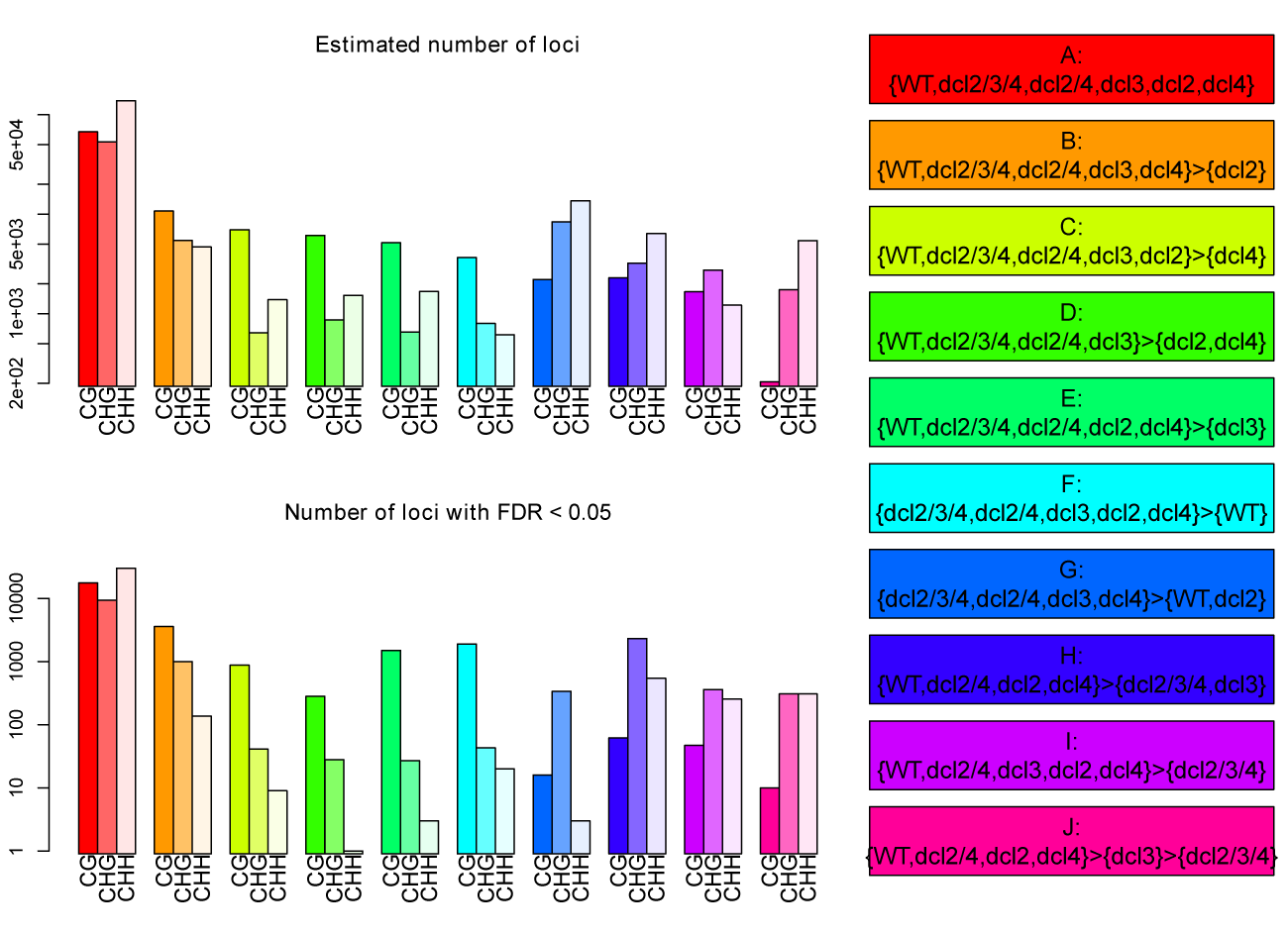
Numbers of loci associated with differential methylation. Expected numbers (log-scale) of methylation loci and the number of loci (log-scale) which can be identified controlling FDR < 0.05 for each of ten patterns of differential methylation, in each of CpG, CHG and CHH contexts.

**Table 1.** Numbers of loci associated with models of differential methylation.

| Model ID | Model definition | Estimated number of loci | Number of identified loci (FDR < 0.05) |
| --- | --- | --- | --- |
| A | {WT,dcl2/3/4,dcl2/4,dcl3,dcl2,dcl4} | CpG: 67365<br>CHG: 53313<br>CHH: 138593 | CpG: 17607<br>CHG: 9310<br>CHH: 30058 |
| B | {WT,dcl2/3/4,dcl2/4,dcl3,dcl4}>{dcl2} | CpG: 10769<br>CHG: 5459<br>CHH: 4686 | CpG: 3587<br>CHG: 996<br>CHH: 137 |
| C | {WT,dcl2/3/4,dcl2/4,dcl3,dcl2}>{dcl4} | CpG: 6989<br>CHG: 644<br>CHH: 1383 | CpG: 877<br>CHG: 41<br>CHH: 9 |
| D | {WT,dcl2/3/4,dcl2/4,dcl3}>{dcl2,dcl4} | CpG: 6131<br>CHG: 866<br>CHH: 1527 | CpG: 281<br>CHG: 28<br>CHH: 1 |
| E | {WT,dcl2/3/4,dcl2/4,dcl2,dcl4}>{dcl3} | CpG: 5167<br>CHG: 654<br>CHH: 1672 | CpG: 1499<br>CHG: 27<br>CHH: 3 |
| F | {dcl2/3/4,dcl2/4,dcl3,dcl2,dcl4}>{WT} | CpG: 3665<br>CHG: 797<br>CHH: 614 | CpG: 1890<br>CHG: 43<br>CHH: 20 |
| G | {dcl2/3/4,dcl2/4,dcl3,dcl4}>{WT,dcl2} | CpG: 2208<br>CHG: 8384<br>CHH: 13657 | CpG: 16<br>CHG: 337<br>CHH: 3 |
| H | {WT,dcl2/4,dcl2,dcl4}>{dcl2/3/4,dcl3} | CpG: 2298<br>CHG: 3220<br>CHH: 6387 | CpG: 62<br>CHG: 2314<br>CHH: 542 |
| I | {WT,dcl2/4,dcl3,dcl2,dcl4}>{dcl2/3/4} | CpG: 1661<br>CHG: 2745<br>CHH: 1219 | CpG: 47<br>CHG: 360<br>CHH: 255 |
| J | {WT,dcl2/4,dcl2,dcl4}>{dcl3}>{dcl2/3/4} | CpG: 207<br>CHG: 1746<br>CHH: 5435 | CpG: 10<br>CHG: 308<br>CHH: 307 |

The ten models selected for further consideration can be roughly partitioned into five classes based on their definitions and the contexts in which they are most commonly found. Model A represents those methylation loci which show no differential methylation. Unsurprisingly, these are common in all contexts of methylation, as the DCL-dependent methylation makes up a small proportion of the total methylation on the *Arabidopsis* genome. The next class is of the models B, C, D & E, describing loci that show some loss of methylation in one or more of the single *dcl* mutants, but not in either the double *dcl2/4* or triple *dcl2/3/4* mutants. These loci are predominantly found in the CpG context, and are particularly rare in CHH context methylation. Model F is also predominantly found in CpG context methylation, and describes loci which show a gain in methylation in all *dcl* mutants relative to the wild-type samples. Similarly, Model G represents a gain in methylation in the majority of the *dcl* mutants over wild-type and the *dcl2* mutant, and is found with confidence only in the CHG methylation context. Models H, I & J represent the canonical changes in sRNA-linked methylation [22], in which there is loss of methylation in either *dcl3* and *dcl2/3/4* relative to wild-type, with Model I somewhat exceptional in that it does not represent a loss of methylation in the *dcl3* mutant but only in the *dcl2/3/4* triple mutant. These loci are predominantly found in CHG and CHH contexts, conforming to the expectation that DCL3 is particularly relevant to the CHG and CHH methylation pathways.

The level of change in methylation varies considerably between models and contexts (Figure 3). For example, the average loss of methylation specific to the *dcl2* mutant (model B) in the CpG mutant is substantial, whereas that specific to the *dcl4* mutant (model C) is much lower (though still detectable at large numbers of loci). Gain in methylation in some or all of the *dcl* mutants can also be substantial (model F; all contexts) or marginal (model G). For CpG context methylation, several of the more significant changes in methylation occur in loci with a short average width (Supplemental Figure 1), notably those in models B, E & F, though this does not necessarily negate their biological significance [23].

**Figure 3.**
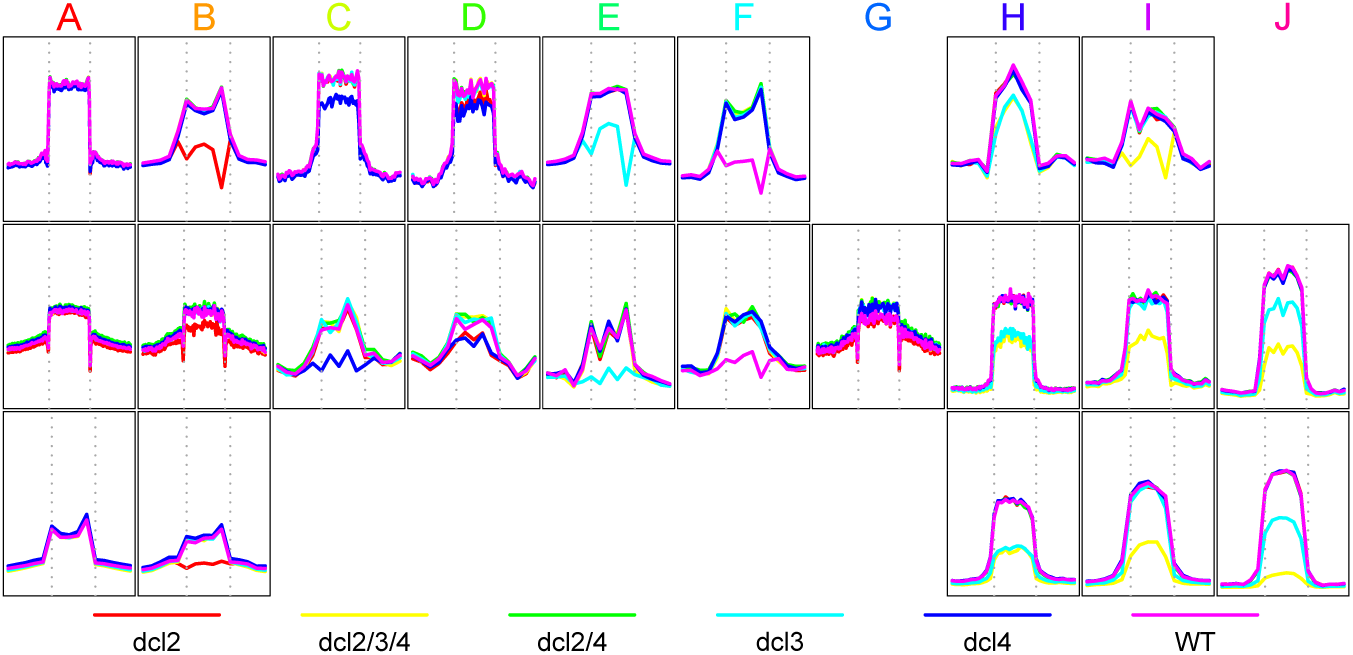
Profiles of cytosine methylation in the dcl mutants for identified differentially methylated loci. Average methylation profiles across the methylation loci (and the surrounding 4Kb) identified for each model/context with an FDR of 5%. Profiles are shown for those model/context combinations in which at least 20 loci at this FDR could be identified.

Some evidence for the biological relevance of the identified classes can be acquired by examining the average methylation profiles for these loci across a range of additional mutants from the Stroud et al [11] dataset (Figure 4). In the CpG context, loci representing models B, C & D, in which methylation is lost in the *dcl2*, *dcl4* or *dcl2/4* mutants also show a substantial average loss of methylation in the *met1* heterozygous mutant, while this effect is much reduced in the non-differential loci (model A) and those loci showing a loss of methylation in the *dcl3* mutant (model E). This effect appears even stronger in loci showing a gain in methylation in all *dcl* mutants over wildtype. Conversely, loci representing models C & D show a reduced loss of CpG methylation in the *ddm1* mutant, perhaps implying a partial independence of these loci from the chromatin remodelling methylation pathway[24].

**Figure 4.**
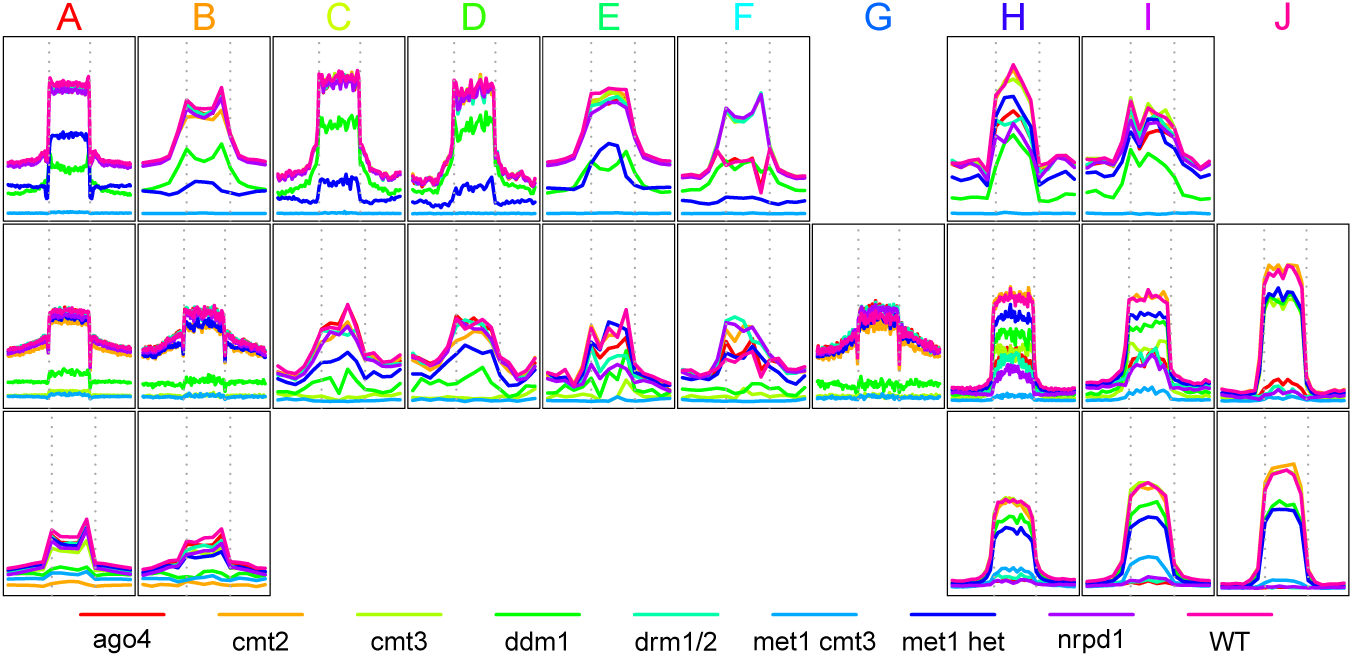
Profiles of cytosine methylation in additional mutants for identified differentially methylated loci. Average methylation profiles in the *ago4, cmt2, cmt3, ddml, drm1/2, metl cmt3, metl het, nrpdl*, and wild-type (WT) samples from Stroud *et al*, across the methylation loci (and the surrounding 4Kb) identified for each model/context with an FDR of 5%. Profiles are shown for those model/context combinations in which at least 20 loci at this FDR could be identified.

In the CHG context, it is notable that the loci showing small gains in average methylation observed in all *dcl* mutants except *dcl2* over wild-type (model G) show a similar gain in the *ago4* and *nrpdl* mutants, supporting the role of the sRNA pathways in repressing methylation at these loci. Also of note is the relative independence of methylation from CMT3 in the loci representing model J, coupled with an increased dependendence on DRM1/2. This suggests a refinement of the model for redundant maintainence of CHG methylation by DRM and CMT3 proposed in Cao *et al* [25] as it indicates that for some RdDM loci it is DRM that is primarily required, and that this correlates with specific patterns of differential behaviour between *dcl3* and *dcl2/3/4.* In the CHH context, perhaps the most notable feature is the partial maintainence of methylation in the *met1/cmt3* mutant in the RdDM loci (models H, I & J). Notably, the average methylation across these loci in the *met1/cmt3* mutant is greatest in those loci affected only in *dcl2/3/4* and not *dcl3* (model I), perhaps indicating that methylation at these loci is more strongly regulated at the establishment phase by 21/22-nt sRNAs [22].

Variation is also marked in the genomic localisation of these models (Figure 5). Loci representing models B, C & D, in which methylation is reduced in either or both of the *dcl2* and *dcl4* mutants, but neither of the double *(dcl2/4*) or triple *(dcl2/3/4*) mutants are found ubiquitously across the genome in the CpG context but are heavily centromeric in CHG and CHH contexts. Conversely, those loci in which methylation is reduced only in the *dcl3* mutant is centromeric in the CpG context. Gains in methylation in some or all of the *dcl* mutants appear evenly distributed across the genome in the CpG context but are strongly centromeric in the CHG context.

**Figure 5.**
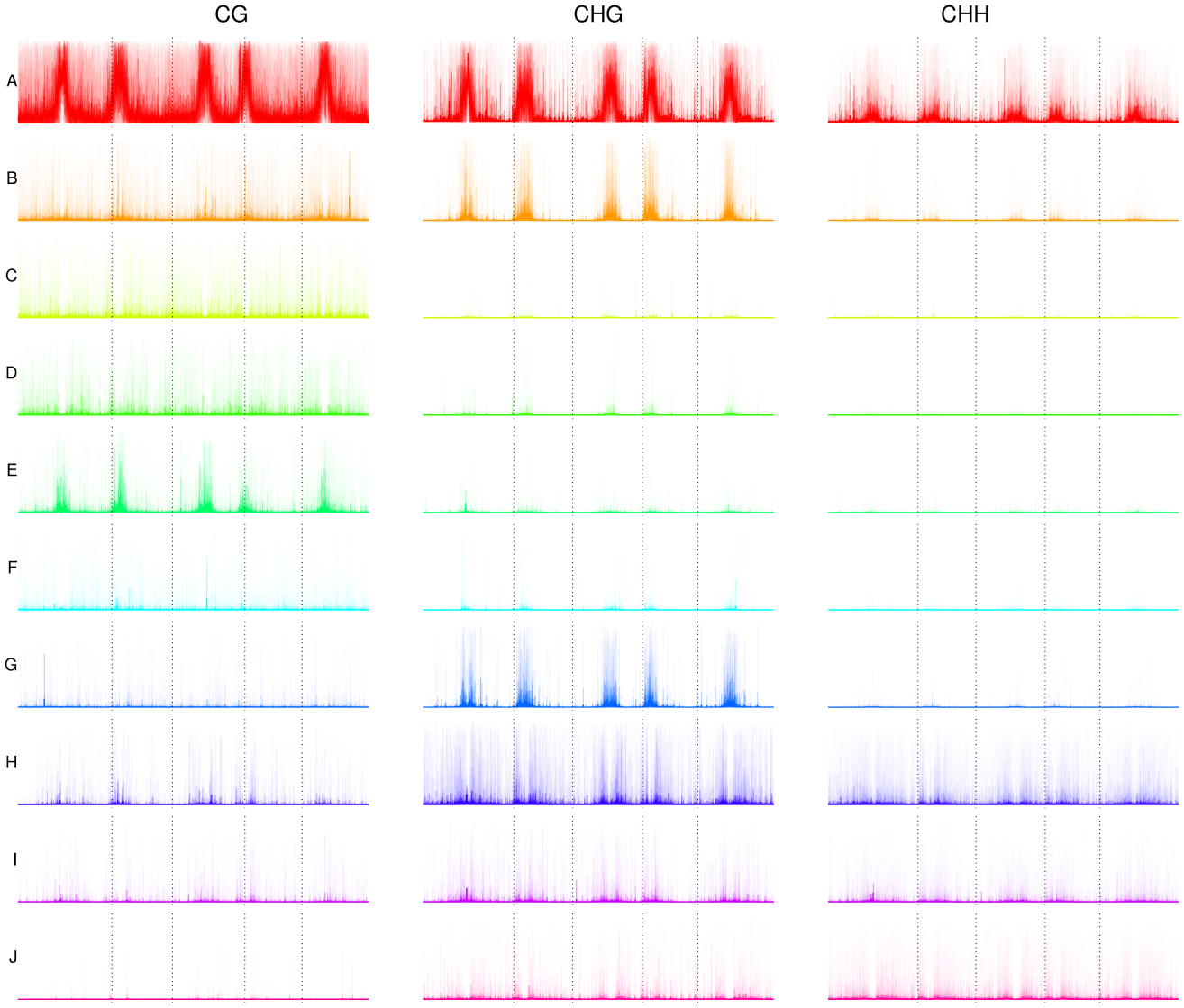
Profiles of model abundance across the genome. Model localisations across the genome; the average posterior likelihood observed at each base within travelling windows of 101, 1001, 10001 and 100001 bases is computed and plotted for each model and methylation context. The transparency of the plotted value increases with window size such that the larger windows have a reduced effect on the plotted values.

### Simulated data

We next compare the performance of the approach developed here to several existing methods for detection of differential methylation using simulation data generated by WGBSSuite [17]. This tool simulates differentially methylated regions based on a complex parameter set allowing a variety of methylation types to be generated. We simulate data based on two basic sets of parameters, one derived from an analysis of CpG methylation in *Arabidopsis* and one desgined to mimic methylation in animal systems (see Methods for parameter details). Using these basic parameters, we evaluate the performance of each method, varying coverage, number of replicates, the magnitude of methylation difference between replicates.

We modified the standard WGBSSuite analysis by including the effects of sample specific non-conversion rates to the data. Non-conversion rates were estimated from the wild-type and various dcl-mutants in the Stroud *et* al[11] dataset, as above. Parameters for a beta distribution approximating the distribution of observed rates were estimated by maximum likelihood. These parameters were then used to simulate non-conversion rates for each sample in a simulation.

The simulated data are evaluated using BSmooth/bsseq [26], MethylKit [27], MethylSig [28] and MethPipe [29]. BSmooth, MethylKit, and MethylSig are implemented as in the WGBSSuite benchmarking, as is a Fisher exact test. These methods primarily rely on the detection of differential methylation at the cytosine level, and construct DMRs from the identified differentially methylated cytosines. MethPipe offers two different implementations, the first similarly based on scores constructed at each cytosine supported by a two-state hidden Markov model used to identify regions of methylation (MethPipe-1), while the second uses a betabinomial regression on the observed data and is recommended for larger sample sets (MethPipe-2).

Performance of the methods is evaluated primarily by constructing a ranked list of DMRs based on each method’s test statistic. As in WGBSSuite’s benchmarking, true postives are defined as the number of truly differentially methylated cytosines within identified DMRs, while false positives are the non-differentially methylated cytosines within the identified DMRs. Figure 6 shows a comparison between the methods for data simulated using parameters intended to produce data similar to those observed in CpG methylation in plant systems. Analyses are carried out using 1, 3, and 10 replicates, and with changes in the proportion of methylated cytosines between experimental groups of 0.05, 0.25 and 0.85.

**Figure 6.**
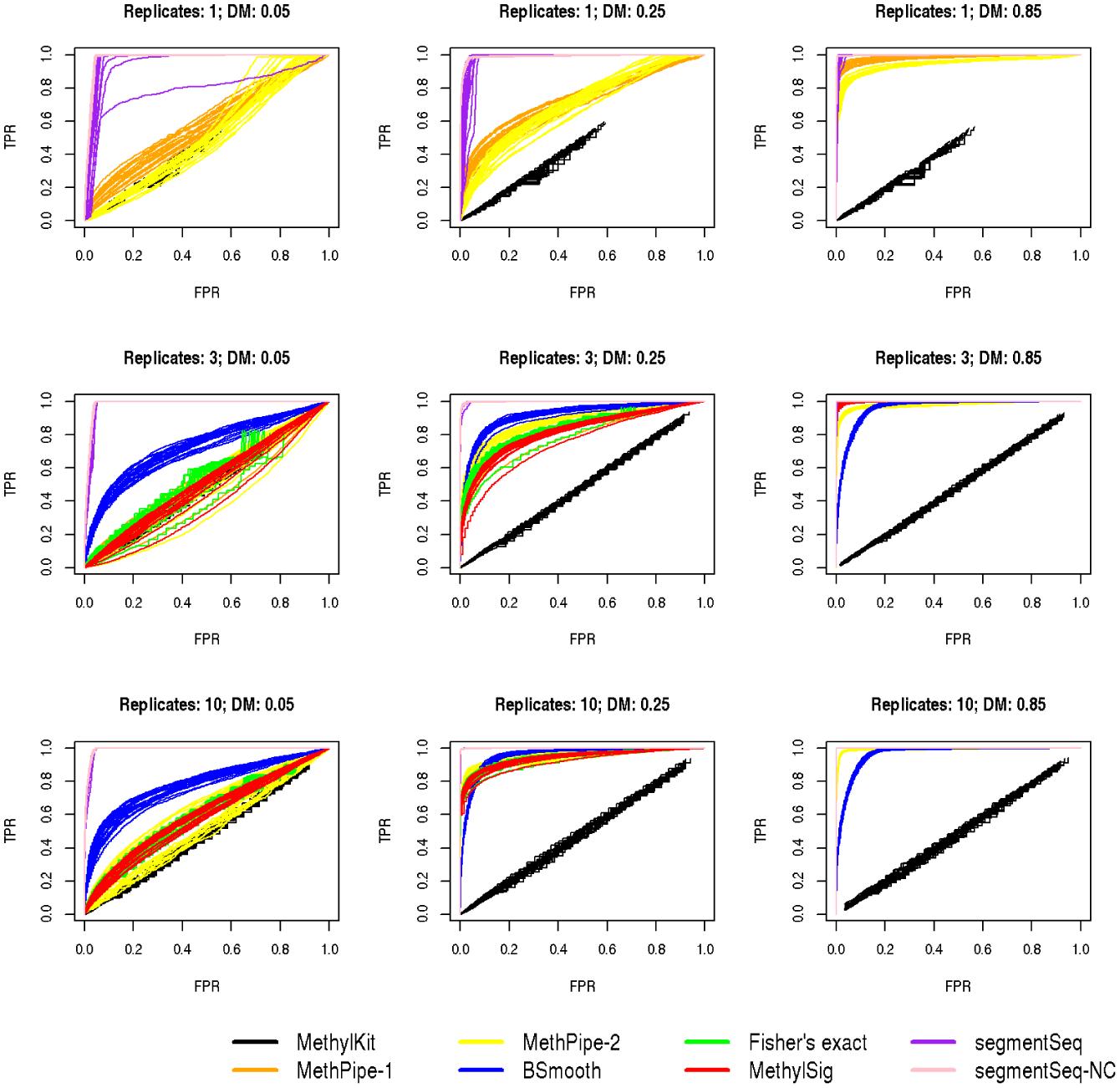
ROC curves from simulated data. ROC curves from simulated data based on WGBSSuite analyses of CpG methylation in *Arabidopsis.* Analyses for 1, 3 and 10 replicates are shown, and for small (0.05), moderate (0.25) and large (0.85) changes in methylation in the differentially methylated regions. Twenty simulations were carried out for each choice of replicate number/difference in methylation and curves for each simulation are shown here.

In all simulations, the segmentSeq/baySeq methods described here perform as well or better than the other methods considered as assessed by the ROC curves. The segmentSeq approach failing to account for non-conversion on average performs well, but shows greater variation and some loss of performance compared to the segmentSeq-NC method which incorporates adjustments for non-conversion. This is particularly true for the experiments with few replicates; with higher numbers of replicates the effect of non-conversion will tend to average out across samples.

For large differences (a proportion shift of 0.85) in methylation all methods are able to detect differentially methylated cytosines in three and ten sample cases with almost perfect accuracy, with the exception of BSmooth and MethylKit. BSmooth shows reduced performance compared to other methods in the ten sample case and MethylKit is unable to make valid calls in any analysis. This is likely due to the design of MethylKit; it is primarily intended for the analysis of reduced representation bisulphit sequencing (RRBS) and does not appear suitable for the substantially lower coverage used in these simulations. For smaller shifts in methylation proportion the increase in performance through the segmentSeq/baySeq approach is more dramatic; this is to be expected as the increased data available to analyse methylation loci rather than individual cytosines gives greater power to detect small differences in methylation. In the analysis without replicates, the segmentSeq/baySeq approach shows substantially better performance over MethPipe and BSmooth for low and moderate differential methylation, with the MethPipe-1 analysis approaching this performance in the high differential case.

## Discussion

A number of methods have previously been developed to analyse high-throughput sequencing of the methylome [7]. These are predominantly focused on the identification of differential methylation and the discovery of differentially methylated regions from grouping differentially methylated cytosines. The methods presented here adopt an alternative strategy in which first methylated and un-methylated regions are identified, and differential methylation is subsequently evaluated. Comparisons on simulated data show that the approach developed here offers substantially more power to detect small changes in methylation across a region when compared to existing methods which operate on a cytosine-by-cytosine scale, without any loss of power in the detection of large shifts in methylation. Accounting for non-conversion rates, where possible, gives a small but consistent improvement in performance, particularly when replication or the level of change in methylation is low. This is perhaps of particular importance in analysing plant methylomes, in which wild-type levels of CHG and CHH context methylation are expected to be low, and consequently loss of methylation is marked by only a small shift in the observed data.

We demonstrate these methods on a subset of the [11] dataset describing the *Arabidopsis* methylome. A primary strength of the approach presented here is its ability to analyse complex relationships between multiple replicate groups. We demonstrate this by the simultaneous analysis of all the *dcl* mutants, together with the wild-type samples contained in Stroud2013. Several novel patterns of methylation are identified through this analysis; most particularly a set of over a thousand CpG loci which lose methylation in the *dcl2* mutant but not the *dcl2/4* double mutant; at somewhat fewer loci we identify similar patterns in CHG and CHH contexts. Similarly, we identify loci which show a reduction in methylation in the *dcl4* mutant but not the *dcl2/4* double mutant, and loci which show a reduction in the *dcl3* but not the *dcl2/3/4* triple mutant. The mechanisms associated with these loci are not directly explicable from the data but it seems likely that there is an antagonistic relationship between the DCL-proteins at at least some of the loci, as previously noted by Bouche *et al* [30]. Support for these loci as biologically meaningful is demonstrated through comparisons with additional mutants of the methylome regulation pathways and through analysis of the genome localisation of the discovered loci.

## Conclusions

The methods described here allow for the identification of methylation loci from multiple sequencing data, the estimation of likelihoods, for each replicate group, that a region is truly methylated above background levels, and ultimately the detection of differential methylated regions. These methods offer a number of significant performance advantages over existing methods for detection of differential methylation, particularly in the detection of small changes in methylation levels and in experiments with low numbers of replicates. These methods also allow for the analysis of complex experimental designs, as demonstrated on a reanalysis of methylation in a set of *dcl* mutants. This analysis demonstrates the potential utility of this method in identifying a variety of methylation loci demonstrating novel interactions between regulatory mechanisms of methylation.

The methods are implemented and released within the `segmentSeq` [9] and `baySeq` [10], available on Bioconductor (http://www.bioconductor.org) [20]. In addition to usability and maintainence advantages, this ensures compatibility with the analyses of sRNA-seq, mRNA-seq *et cetera* already developed in these packages. Results acquired by high-throughput sequencing of methylation can thus be readily integrated with these other -omic data, allowing the differential methylome to be incorporated into in systems level analyses.

## Methods

### Candidate loci and nulls

We begin our analysis of these data by defining a set of *candidate loci* which may plausibly represent some methylation loci. A candidate locus begins and ends at some cytosine with a minimal proportion *p_min_* of methylation in at least one sequencing library. Considering all such loci is computationally infeasible and so filters are required to exclude implausible candidates and reduce the computational effort required. If two cytosines with a proportion of methylation above pmin are within some minimal distance *d_min_* they are assumed to lie within the same locus. We further restrict the set by removing from consideration any candidate locus containing a region greater than *λmax* that contains no cytosine with a proportion of methylation above *p_min_*. Candidate loci may be defined with respect to a single strand (by default), or combine the observed data from both strands.

We define the set of *candidate nulls*, regions which may represent a region without significant methylation, by considering the gaps between candidate loci. We refer to the regions separating each candidate locus from its nearest neighbour (in either direction) as ‘empty’. Candidate nulls consist of the union of the set of ‘empty’ regions, the set of candidate loci extended into the empty region to their left, the set of candidate loci extended into the empty region to their right, and the set of candidate loci extended into the empty regions to both the left and right.

### Classification of candidate loci

The data pertaining to the candidates defined above are the number of methylated and un-methylated cytosines sequenced and aligning to these loci for each sample. Biological replication is defined in terms of *replicate groups*, non-intersecting sets of biological replicates. Thus the samples may be thought of as the set 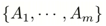 with a replicate structure defined by 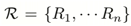 where j ∈ *R_q_* if and only if sample *A_j_* is a member of replicate group *q*. We then identify those candidates which represent at least part of a true locus of methylation, given the observed data for each replicate group

For a replicate group *R_q_* and segment *i* we consider the total number of methylated and unmethylated cytosines 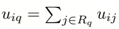 and 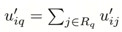 respectively. We assume that these data are described by a binomial distribution with parameter *p_iq_* which has a beta prior distribution with parameters (*α,β*); we use an uninformative Jeffreys prior of 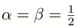. The posterior distribution of the parameter piq is then a beta distribution with parameters 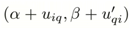 A segment is identified as a methylation ‘locus’ if the posterior likelihood that *p_qj_* > *t* exceeds some critical value. Similarly, we can classify candidate nulls as true representatives of a null region by identifying those candidates with a posterior likelihood that *p_qj_* < *t* exceeding some critical value. By default, we use a required likelihood of 90%. The parameter t is intended to provide a threshold distinguishing regions of ‘true’ or biologically relevant methylation from low level background methylation attributable to biological or technical noise. The appropriate value for this parameter is contingent on many factors, including organism, methylation context, heterogeneity of sample and the assignment of biological meaning; we use a value of 20% here across all analyses with the caveat that this may be more or less appropriate to any individual experiment.

The above analysis neglects the effect of non-conversion rates on the observed values for *u_qj_* and *u’_qj_*. We can find no closed form expression for the posterior if the effects of non-conversion rates on the distribution of the data are accounted for. However, we can normalise the observed data by the expected non-conversion rates by setting 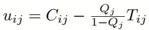 and 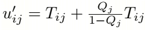, where *Q_j_* is the nonconversion rate for sample *j*.

### Consensus loci

Given a classification on the set of candidate loci and nulls, we identify a set of consensus loci given the classifications on sets of overlapping candidates in a similar manner to that described for siRNA loci [31]. We begin by assuming that a true locus of methylation should not contain a null region within a replicate group in which the locus is methylated. Thus, if some candidate locus *l_r_* is classified as a locus in replicate groups Ψ*_r_*, and there exists some candidate null *n_s_* that lies completely within *l_r_* and is classified as a null in one or more of the replicate groups Ψ*r*, we discard the locus *l_r_*. Of the remaining candidate loci, we then rank those that remain by the number of replicate groups in which they are classified as a locus, settling ties by considering the longer candidate locus. The consensus loci are then formed by choosing all those candidate loci that do not overlap with some higher ranked candidate, giving a non-overlapping set of loci on each strand. ‘Null’ loci are defined as the contiguous regions of the genome containing no identified locus.

### Likelihood of data

We can compute posterior likelihoods of methylation and differential methylation on the identified loci through application of the empirical Bayesian methods described in Hardcastle[10]. We achieve this by defining a distribution on the data accounting for biological variation between replicates. Ignoring issues of non-conversion, we would assume that the data in equivalently methylated samples are beta-binomially distributed as in a straightforward analysis of paired data[31].

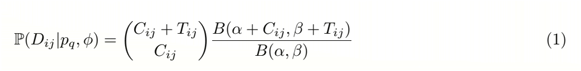

Eqn 1 defines the likelihood of the observed data *D_ij_*, consisting of the observed number of methylated *(C_ij_*) and unmethylated (*T_ij_*) sequenced cytosines at the *i*th locus and *j*th sample, given a proportion of methylation *p* and a dispersion parameter Φ capturing the level of variation between biological replicates. Then 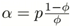, 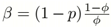. Following our previous work[10], a joint distribution on {*p, Φ*} may be empirically estimated from the observed data.

If non-conversion rates are estimable, we can first normalise the observed data as above and proceed assuming a beta-binomial distribution. However, this does not fully account for the stocasticity of non-conversion events at each cytosine. A full analysis incorporating non-conversion events requires that the data within each sample *j* are assumed to be the sum of a binomial distribution with success parameter *Q_j_* (the rate of non-conversion) and a beta-binomial distribution with parameters *p* (the expected proportion of methylated cytosines) and dispersion parameter Φ. Then the likelihood of the observed data *D_jk_* at a single locus *i* for a sample *j* is given by

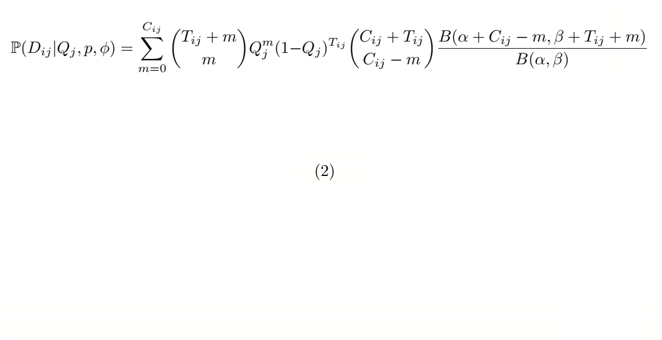

where *m* is the number of unconverted unmethylated cytosines and the remaining parameters are as in Eqn 1.

### Posterior Likelihoods of Methylation

We can estimate posterior likelihoods of methylation for each replicate group and locus using the methods described in Hardcastle[10]. For a sampled locus, we estimate by maximum likelihood methods for each replicate group q the parameters {*p, Φ*}, in which the dispersion parameter Φ is assumed to be preserved across replicate groups and *p* is not. By repeating (without replacement) the sampling of loci, we build an empirical joint distribution on the parameters for the methylation of loci within each replicate group. We similarly derive an empirical joint distribution on for null regions. Given these distributions, we are able to calculate posterior likelihoods of methylation for each locus and replicate group. Regions exhibiting various patterns of differential methylation can be similarly identified using the density function defined in Eqn 2 or Eqn 1 (neglecting non-conversion rates) in the `baySeq R` package.

Since the definition of differential methylation is primarily concerned with a shift in ratios between the number of methylated cytosines and the number of unmethylated cytosines, it is possible for long regions of low methylation to exhibit patterns of differential methylation. We thus find improved performance by combining the likelihood of differential methylation within a locus with the likelihood of that locus being methylated in at least one replicate group. The final statistic used to identify DMRs with this method is thus:

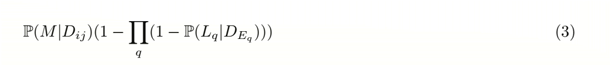

where 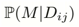 is the likelihood of a model *M* of differential methylation given the observed data in each sample at the ith defined region, and 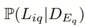 is the likelihood that the *i*th region defines an expressed locus within replicate group *q*.

### Simulation studies

Data are simulated using WGBSSuite version 0.4, modified to include the effect of non-conversion rates by introducing a binomially distributed error in the number of methylated cytosines reported at each site. Plant-representative parameters were estimated by applying the `analyse_WGBS.R` script provided by WGBSSuite to the first million CpG cytosines on chromosome 1 of a single wild-type sample and the *dcl2/3/4* triple mutant in the Stroud *et* al[11] dataset. Data were then simulated for one hundred thousand cytosines using the following command, where XXX and YYY are placeholders for the number of samples in each replicate group and the magnitude of difference in the proportion of methylation respectively.

Rscript simulate_WGBS.R 100000 0.851721358101034 0.0158503393648432 +0.2 0.2 14.064221221 14.064221221 XXX 2 YYY 0.5 +0.0267845702643185,0.00339225464276114 plant_simulation binomial +0.237414948538599,1e-15 0.0577772070946612,0.34

Version 0.4.4 of methylSig, 1.14 of methylKit, 1.10.0 of BSmooth and 3.4.2 of MethPipe were used to analyse these data using default settings for all libraries. Versions 2.9.5 of segmentSeq and 2.8.0 of baySeq were used to implement the approach described here.

## Competing interests

The authors declare that they have no competing interests.

## Acknowledgements

This work was supported by European Research Council Advanced Investigator Grant ERC-2013-AdG 340642 TRIBE.

## References

1. Zhang, X., Yazaki, J., Sundaresan, A., Cokus, S., Chan, S.W.-L., Chen, H., Henderson, I.R., Shinn, P., Pellegrini, M., Jacobsen, S.E., Ecker, J.R.: Genome-wide high-resolution mapping and functional analysis of DNA methylation in arabidopsis. Cell 126(6), 1189–201 (2006). doi:10.1016/j.cell.2006.08.003

2. Berdasco, M., Alcázar, R., García-Ortiz, M.V., Ballestar, E., Fernández, A.F., Roldán-Arjona, T., Tiburcio, A.F., Altabella, T., Buisine, N., Quesneville, H., Baudry, A., Lepiniec, L., Alaminos, M., Rodríguez, R., Lloyd, A., Colot, V., Bender, J., Canal, M.J., Esteller, M., Fraga, M.F.: Promoter DNA hypermethylation and gene repression in undifferentiated Arabidopsis cells. PloS One 3(10), 3306 (2008). doi:10.1371/journal.pone.0003306

3. Tsukahara, S., Kobayashi, A., Kawabe, A., Mathieu, O., Miura, A., Kakutani, T.: Bursts of retrotransposition reproduced in Arabidopsis. Nature 461(September),3–7 (2009). doi:10.1038/nature08351

4. Clark, S.J., Statham, A., Stirzaker, C., Molloy, P.L., Frommer, M.: DNA methylation: bisulphite modification and analysis. Nature Protocols 1(5), 2353–64 (2006). doi:10.1038/nprot.2006.324

5. Bock, C.: Analysing and interpreting DNA methylation data. Nature Reviews Genetics 13(10), 705–19 (2012). doi:10.1038/nrg3273

6. Hardcastle, T.J.: High-throughput sequencing of cytosine methylation in plant DNA. Plant Methods 9(1), 16 (2013)

7. Robinson, M.D., Kahraman, A., Law, C.W., Lindsay, H., Nowicka, M., Weber, L.M., Zhou, X.: Statistical methods for detecting differentially methylated loci and regions. Frontiers in genetics 5, 324 (2014). doi:10.3389/fgene.2014.00324

8. Zhao, J.-H., Fang, Y.-Y., Duan, C.-G., Fang, R.-X., Ding, S.-W., Guo, H.-S., Bird, A., Chan, S.W., Henderson, I.R., Jacobsen, S.E., Zhang, X., Lister, R., Lindroth, A.M., Cao, X., Jacobsen, S.E., Matzke, M., Kanno, T., Daxinger, L., Huettel, B., Matzke, A.J., Zhang, H., Zhu, J.-K., Dalakouras, A., Wassenegger, M., Zhao, M., Leon, D.S., Delgadillo, M.O., Garcia, J.A., Simon-Mateo, C., Dalakouras, A., Dadami, E., Zwiebel, M., Krczal, G., Wassenegger, M., Wierzbicki, A.T., Haag, J.R., Pikaard, C.S., Law, J.A., Jacobsen, S.E., Wu, L., Mao, L., Qi, Y., Nuthikattu, S., Stroud, H., Greenberg, M.V., Feng, S., Bernatavichute, Y.V., Jacobsen, S.E., Lee, T.F., Wang, H., Hao, L., Shung, C.Y., Sunter, G., Bisaro, D.M., Wang, H., Buckley, K.J., Yang, X., Buchmann, R.C., Bisaro, D.M., Buchmann, R.C., Asad, S., Wolf, J.N., Mohannath, G., Zhang, Z., Yang, X., Ivanov, K.I., Canizares, M.C., Li, H.W., Guo, H.S., Ding, S.W., Gonzalez, I., Duan, C.G., Hamera, S., Song, X., Su, L., Chen, X., Fang, R., Feng, L., Duan, C.-G., Guo, H.-S., Cokus, S.J., Li, Y., Wang, H., Mi, S., Takeda, A., Iwasaki, S., Watanabe, T., Utsumi, M., Watanabe, Y., Girard, A., Hannon, G.J., Sarazin, A., Voinnet, O., Creasey, K.M., Zhang, X., Zhong, X., Mosher, R.A., Schwach, F., Studholme, D., Baulcombe, D.C., Mirouze, M., Mari-Ordonez, A., Slotkin, R.K., Han, B.W., Wang, W., Li, C., Weng, Z., Zamore, P.D., Mohn, F., Handler, D., Brennecke, J., Siomi, H., Siomi, M.C., Calvi, B.R., Gelbart, W.M., Dupressoir, A., Heidmann, T., Pasyukova, E., Nuzhdin, S., Li, W., Flavell, A.J., Ostertag, E.M., Lau, N.C., Brennecke, J., Brower-Toland, B., Carmell, M.A., Zahid, K., Allen, G.C., Flores-Vergara, M.A., Krasynanski, S., Kumar, S., Thompson, W.F., Li, R., Pathak, R.R., Lochab, S., Mortazavi, A., Williams, B.A., McCue, K., Schaeffer, L., Wold, B., Kim, K.I., van de Wiel, M.A.: Genome-wide identification of endogenous RNA-directed DNA methylation loci associated with abundant 21-nucleotide siRNAs in Arabidopsis. Scientific Reports 6, 36247 (2016). doi:10.1038/srep36247

9. Hardcastle, T.J., Kelly, K.A., Baulcombe, D.C.: Identifying small interfering RNA loci from high-throughput sequencing data. Bioinformatics 28(4), 457–463 (2012). doi:10.1093/bioinformatics/btr687

10. Hardcastle, T.J.: Generalized empirical Bayesian methods for discovery of differential data in high-throughput biology. Bioinformatics 32(2), 195–202 (2016). doi:10.1093/bioinformatics/btv569

11. Stroud, H., Greenberg, M.C., Feng, S., Bernatavichute, Y., Jacobsen, S.: Comprehensive Analysis of Silencing Mutants Reveals Complex Regulation of the Arabidopsis Methylome. Cell 152(1), 352–364 (2013). doi:10.1016/j.cell.2012.10.054

12. Chapman, E.J., Carrington, J.C.: Specialization and evolution of endogenous small RNA pathways. Nature Reviews Genetics 8(11), 884–896 (2007). doi:10.1038/nrg2179

13. Gasciolli, V., Mallory, A.C., Bartel, D.P., Vaucheret, H.: Partially Redundant Functions of Arabidopsis DICER-like Enzymes and a Role for DCL4 in Producing trans-Acting siRNAs. Current Biology 15(16), 1494–1500 (2005). doi:10.1016/j.cub.2005.07.024

14. Panda, K., Slotkin, R.K.: Proposed mechanism for the initiation of transposable element silencing by the RDR6-directed DNA methylation pathway. Plant Signaling & Behavior 8(8) (2013). doi:10.4161/psb.25206

15. Bond, D.M., Baulcombe, D.C.: Epigenetic transitions leading to heritable, RNA-mediated de novo silencing in Arabidopsis thaliana. Proceedings of the National Academy of Sciences 112(3), 917–922 (2015). doi:10.1073/pnas.1413053112

16. Matzke, M.A., Kanno, T., Matzke, A.J.M.: RNA-Directed DNA Methylation: The Evolution of a Complex Epigenetic Pathway in Flowering Plants. Annual Review of Plant Biology 66(1), 243–267 (2015). doi:10.1146/annurev-arplant-043014-114633

17. Rackham, O.J.L., Dellaportas, P., Petretto, E., Bottolo, L.: WGBSSuite: simulating whole-genome bisulphite sequencing data and benchmarking differential DNA methylation analysis tools. Bioinformatics 31(14), 2371–3 (2015). doi:10.1093/bioinformatics/btv114

18. Team, R.D.C.: R: A Language and Environment for Statistical Computing. Vienna, Austria (2012). http://www.r-project.org

19. Hardcastle, T.J., Kelly, K.A.: baySeq: empirical Bayesian methods for identifying differential expression in sequence count data. BMC Bioinformatics 11(1), 422 (2010). doi:10.1186/1471-2105-11-422

20. Gentleman, R.C., Carey, V.J., Bates, D.M., Bolstad, B., Dettling, M., Dudoit, S., Ellis, B., Gautier, L., Ge, Y., Gentry, J., Hornik, K., Hothorn, T., Huber, W., Iacus, S., Irizarry, R., Leisch, F., Li, C., Maechler, M., Rossini, A.J., Sawitzki, G., Smith, C., Smyth, G., Tierney, L., Yang, J.Y.H., Zhang, J.: Bioconductor: open software development for computational biology and bioinformatics. Genome Biology 5(10), 80 (2004). doi:10.1186/gb-2004-5-10-r80

21. Krueger, F., Andrews, S.R.: Bismark: a flexible aligner and methylation caller for Bisulfite-Seq applications. Bioinformatics 27(11), 1571–2 (2011). doi:10.1093/bioinformatics/btr167

22. Bond, D.M., Baulcombe, D.C.: Small RNAs and heritable epigenetic variation in plants. Trends in Cell Biology 24(2), 100–107 (2014). doi:10.1016/j.tcb.2013.08.001

23. Xu, J., Pope, S.D., Jazirehi, A.R., Attema, J.L., Papathanasiou, P., Watts, J.A., Zaret, K.S., Weissman, I.L., Smale, S.T.: Pioneer factor interactions and unmethylated CpG dinucleotides mark silent tissue-specific enhancers in embryonic stem cells. Proceedings of the National Academy of Sciences of the United States of America 104(30), 12377–82 (2007). doi:10.1073/pnas.0704579104

24. Zemach, A., Kim, M.-Y., Hsieh, P.-H., Coleman-Derr, D., Eshed-Williams, L., Thao, K., Harmer, S., Zilberman, D.: The Arabidopsis Nucleosome Remodeler DDM1 Allows DNA Methyltransferases to Access H1-Containing Heterochromatin. Cell 153(1), 193–205 (2013). doi:10.1016/j.cell.2013.02.033

25. Cao, X., Aufsatz, W., Zilberman, D., Mette, M.F., Huang, M.S., Matzke, M., Jacobsen, S.E.: Role of the DRM and CMT3 methyltransferases in RNA-directed DNA methylation. Current Biology 13(24), 2212–7 (2003)

26. Hansen, K.D., Langmead, B., Irizarry, R.A.: BSmooth: from whole genome bisulfite sequencing reads to differentially methylated regions. Genome Biology 13(10), 83 (2012). doi:10.1186/gb-2012-13-10-r83

27. Akalin, A., Kormaksson, M., Li, S., Garrett-Bakelman, F.E., Figueroa, M.E., Melnick, A., Mason, C.E.: methylKit: a comprehensive R package for the analysis of genome-wide DNA methylation profiles. Genome Biology 13(10), 87 (2012). doi:10.1186/gb-2012-13-10-r87

28. Park, Y., Figueroa, M.E., Rozek, L.S., Sartor, M.A.: MethylSig: a whole genome DNA methylation analysis pipeline. Bioinformatics 30(17), 2414–22 (2014). doi:10.1093/bioinformatics/btu339

29. Song, Q., Decato, B., Hong, E.E., Zhou, M., Fang, F., Qu, J., Garvin, T., Kessler, M., Zhou, J., Smith, A.D.: A reference methylome database and analysis pipeline to facilitate integrative and comparative epigenomics. PloS one 8(12), 81148 (2013). doi:10.1371/journal.pone.0081148

30. Bouche, N., Lauressergues, D., Gasciolli, V., Vaucheret, H.: An antagonistic function for Arabidopsis DCL2 in development and a new function for DCL4 in generating viral siRNAs. The EMBO Journal 25(14), 3347–3356 (2006). doi:10.1038/sj.emboj.7601217

31. Hardcastle, T.J., Kelly, K.A.: Empirical Bayesian analysis of paired high-throughput sequencing data with a beta-binomial distribution. BMC Bioinformatics 14(1), 135 (2013). doi:10.1186/1471-2105-14-135

